# Ancient mitogenomes reveal evidence for the late Miocene dispersal of mergansers to the Southern Hemisphere

**DOI:** 10.1101/2023.09.03.556105

**Authors:** Nicolas J. Rawlence, Alexander J. F. Verry, Theresa L. Cole, Lara D. Shepherd, Alan J. D. Tennyson, Murray Williams, Jamie R. Wood, Kieren J. Mitchell

## Abstract

Mergansers are riverine and coastal piscivorous ducks that are widespread throughout North America and Eurasia but uncommon in the Southern Hemisphere. One species occurs in South America and at least two extinct species from New Zealand. It has been proposed that Southern Hemisphere mergansers were founded by independent dispersal events from the Northern Hemisphere. However, some morphological and behavioural evidence suggests that Southern Hemisphere mergansers may be closely related to one another. They share several characteristics that differ from Northern Hemisphere mergansers (e.g., non-migratory vs. migratory, sexual monochromatism vs. sexual dichromatism, long vs. short pair bonds). We sequenced complete mitogenomes from the Brazilian merganser and an extinct merganser from New Zealand—the Auckland Island merganser. Our results show the Brazilian and Auckland Island mergansers form a monophyletic clade with the common merganser, and that a sister relationship between Southern Hemisphere mergansers cannot be rejected. We cannot exclude the possibility that the Brazilian and Auckland Island mergansers descend from a single dispersal event from the Northern Hemisphere at least seven million years ago. Nuclear (palaeo)genomic data may help to resolve the relationship between living and extinct mergansers, including merganser fossils from New Zealand that have not been subjected to palaeogenetic analysis.

## INTRODUCTION

The ducks in the tribe Mergini are a group of riverine and seasonally coastal fish-eating ducks that have a widespread Northern Hemisphere distribution but are uncommon in the Southern Hemisphere (Kear, 2005; Williams *et al*. 2012; Williams *et al*. 2014). One group of the Mergini are the mergansers (*Mergus* spp.), which are characterised by a serrated bill, and include the endangered scaly-sided merganser (*M. squamatus* Gould, 1864) from northeast Asia; the common merganser (*M. merganser* Linnaeus, 1758) and red-breasted merganser (*M. serrator* Linnaeus, 1758), which have widespread Northern Hemisphere distributions; the critically endangered Brazilian merganser (*M. octosetaceus* Vieillot, 1817); and two currently recognised extinct species from the New Zealand region—*M. australis* Hombron and Jacquinot, 1841 and *M. milleneri* Williams and Tennyson, 2014 from the Auckland and Chatham Islands, respectively. While the hooded merganser *Lophodytes cucullatus* (Linnaeus, 1758), previously *M. cucullatus*, from North America has a serrated bill, it is not considered a “true” merganser (e.g. Buckner *et al*. 2018; Lavretsky *et al*. 2021). The taxonomic relationship of the smew *Mergellus albellus* (Linnaeus, 1758) from Eurasia is currently unresolved; it is sometimes suggested to be more closely related to *Mergus* and *Lophodytes* or to goldeneyes (*Bucephala* spp.) (Livezey, 1995; Buckner *et al*. 2018; Lavretsky *et al*. 2021).

The now extinct Auckland Island merganser *M. australis* (or miuweka) was formally described in 1841, based on a specimen collected on the sub-Antarctic Auckland Islands, 465 km south of mainland New Zealand. Rare late Holocene-aged merganser bones have been found in coastal sand dune deposits (including Māori middens) on New Zealand’s three main islands (Stewart, North and South), and the Auckland and Chatham Islands (Tennyson & Martinson, 2007; Williams *et al*. 2014; Tennyson, 2020). Bones from the latter were recently described as a distinct species *M. millerneri*, which was smaller than the nominate *M. australis*, with a shorter skull, relatively shorter premaxilla, smaller sternum and keel, relatively shorter wing bones and a narrower pelvis (Williams *et al*., 2014). The taxonomic status of merganser bones from mainland New Zealand is unresolved (i.e. cannot be assigned to either *M. australis* or *M. millerneri*), and are currently recognised as *Mergus* spp. (Checklist Committee, 2022).

Mergansers in the New Zealand region are mainly thought to have occupied a riverine and seasonally coastal habitat (e.g. sheltered bays during winter; Kear, 2005; Williams *et al*., 2012; Williams *et al*. 2014). It is likely that they mainly nested in tree cavities, but also caves in some instances, as the remains of adults, chicks, and eggs have been found within a cave on the Chatham Islands (Williams *et al*., 2014). By the 17^th^ Century, mergansers had been extirpated from the Chatham Islands and mainland New Zealand, and survived only on the Auckland Islands. A combination of subsistence hunting, predation from the Pacific rat (*Rattus exulans*) and Polynesian dog (*Canis familiaris*), led to the extinction of mergansers across most of their prehistoric range (Tennyson & Martinson, 2007; Greig & Rawlence, 2021). On the Auckland Islands, predation from introduced pigs (*Sus scrofa*) and cats (*Felis catus*), and collecting for the museum trade in the 19^th^ Century, resulted in their extinction—indeed the last known Auckland Island merganser specimen was shot and collected in January 1902 (Williams, 2012).

The only extant merganser in the Southern Hemisphere—the critically endangered Brazilian merganser—is one of the rarest birds in the world, comprising only 250 wild individuals. It is split across three remnant populations in Brazil, but once had a more widespread historical distribution encompassing Argentina and Paraguay (Vilaca *et al*., 2012; Maia *et al*., 2020). The Brazilian merganser has undergone a significant population bottleneck, yet different remnant populations can still be genetically identified (Maia *et al*., 2020). Like mergansers from the New Zealand region, the Brazilian merganser occupies riverine habitats, and often nests in tree cavities or rock crevasses (Vilaca *et al*., 2012; Maia *et al*., 2020).

It has been proposed that the Southern Hemisphere mergansers were founded by independent dispersal events to the New Zealand region and South America from the Northern Hemisphere (e.g. Livezey, 1995). Based on behavioural characteristics, Johnsgard (1961) tentatively assigned the Brazilian merganser as sister species to a clade comprising the other *Mergus* species, with the Auckland Island merganser as the sister species of the common merganser and scaly-sided merganser. In contrast, using morphological characters, Livezey (1989, 1995) assigned the Auckland Island merganser, then Brazilian merganser, as successive sister species to all other *Mergus* species, though with weak to moderate bootstrap support. Using mitochondrial DNA (mtDNA) sequences, Buckner *et al*. (2018) showed the Brazilian merganser is the sister species to the scaly-sided merganser (Buckner *et al*., 2018) albeit weakly supported. However, some evidence suggests that Southern Hemisphere mergansers may be more closely related to one another, potentially even sister species, as they share several behavioural (e.g. non-migratory, long pair bonds) and morphological (e.g. sexually monochromatic) characteristics, in contrast to their Northern Hemisphere congeners (e.g. migratory, short pair bonds, sexual dichromatism; Livezey, 1995). Genetic studies of extinct Southern Hemisphere avian species have also recently revealed unexpected evolutionary connections between birds from New Zealand, South America, and Africa (e.g. Mitchell *et al*., 2014a,b; Boast *et al*., 2019; Rawlence *et al*., 2022). As such, the phylogenetic relationships of the Southern Hemisphere mergansers, when their ancestors arrived in the region, and from where, remains unresolved.

In this study, which is the first genetic study on a New Zealand *Mergus* species, we sequenced mitochondrial genomes (mitogenomes) from historical museum specimens from the Auckland Island merganser and Brazilian merganser, and analysed them within a phylogenetic framework of Mergini mitogenomes (Liu *et al*., 2012; Lavretsky *et al*., 2021). These data were used to determine the phylogenetic relationships and divergence dates within mergansers.

## MATERIALS AND METHODS

### Brazilian merganser mitogenome sequencing

A toepad from a historical Brazilian merganser was sourced from the American Museum of Natural History (AMNH-748583). This specimen is a skin of an adult female bird, and was collected by A. Rivas on 18^th^ Aug 1952 from Puerto Piraiu in Misiones, Argentina. DNA extraction, PCR set-up, and DNA library preparation was performed within a purpose-built ancient DNA laboratory at the University of Otago (Otago Palaeogenetics Laboratory), physically isolated from other molecular laboratories, and subject to regular decontamination with bleach and UV irradiation (Knapp *et al*., 2012). Strict ancient DNA procedures were followed to minimise contamination of samples with exogenous DNA. Downstream procedures (e.g. library indexing and hybridisation capture enrichment) were undertaken within a separate modern genetics laboratory. Ancient DNA extraction followed Verry *et al*. (2019).

Double-stranded DNA libraries were prepared from 16 µL of DNA extract following the BEST protocol of Carøe *et al*. (2018), with modifications, as stipulated in Verry *et al*. (2022). A blank library control was also included to monitor for contamination. qPCR was used to determine the optimal number of PCR-cycles for the indexing PCR using the primer pair IS7/IS8 (Meyer & Kircher, 2010) and the qPCR protocol of Gansauge and Meyer (2013). For indexing amplification, each library was divided into 4 x 25 µL reactions each containing 5 µL of DNA library, 1X PCR Buffer II, 2.5 mM MgCl_2_, 0.8 mg/µL BSA, 0.2 µM of each forward and reverse indexing primer (P5 and P7, with each featuring a seven-nucleotide barcode; see Gansauge & Meyer 2013), 0.25 mM dNTPs, and 2 U of Amplitaq Gold DNA Polymerase, made up to 25 µL with double-distilled MilliQ water. The thermocycling profile for indexing PCR consisted of 95°C 5 min; 12-23 cycles of 95°C 30 sec, 60°C 30 sec, 72°C 1 min, followed by 10 min at 72°C. The four separate reactions for each library were pooled and purified using the QIAGEN MinElute PCR purification kit.

DNA libraries were enriched using a commercially synthesised custom myBaits kit (Arbor Biosciences) consisting of biotinylated 80 mer RNA baits designed to capture avian mitochondrial DNA (excluding the control region/D-loop; see Mitchell *et al*., 2014b). This bait set has been used previously to retrieve mitochondrial DNA from a wide range of avian taxa (e.g. Mitchell *et al*. 2016; Verry *et al*., 2022), and was designed using published whole mitogenomes from a variety of taxa including palaeognaths and neoavians (e.g. acanthisittid wrens, galloanseres; see Mitchell *et al*., 2014a, b). DNA-RNA hybridisation capture enrichment for mitochondrial DNA followed the standard myBaits protocol (version 4.01) with the following modifications: baits were diluted by 1:4; the hybridisation period was extended to 48 hours and consisted of 8 hours at 60°C followed by 40 hours at 55°C; all wash steps were conducted at 55°C. Additionally, qPCR was used to determine the number of cycles necessary to re-amplify each library using the qPCR protocol detailed above, except that the IS5 and IS6 reamplification primers were used (see Meyer & Kircher, 2010). Following qPCR, DNA libraries were re-amplified using the same reaction conditions/thermocycling profile as indexing PCR, with the indexing primers replaced by IS5 and IS6 (Meyer and Kircher, 2010). PCRs were pooled by sample and purified using AMPure XP (at a ratio of 1.2:1 AMPure to PCR product to remove DNA sequences less than 150 bp, i.e. adapter dimers) and eluted in 21 uL of elution buffer. DNA libraries were quantified via Qubit and Qiaxcel. Enriched DNA libraries were then diluted to 10 nM, pooled, and sent to the Garvin Medical Institute for sequencing on an Illumina NextSeq 550 sequencing platform using 2 x 75 bp (150 cycle, paired-end) sequencing chemistry.

### Auckland Island merganser mitogenome sequencing

A toepad sample from a historical Auckland Island merganser was sourced from Canterbury Museum (Av1583). This specimen is a skin of an adult male bird collected by Robert A. Wilson on 30^th^ Oct 1891, probably at Waterfall Inlet, Auckland Island (see Williams, 2012 and references therein). DNA was extracted at the dedicated ancient DNA facility at the Long-Term Ecology Laboratory, Manaaki Whenua – Landcare Research (Lincoln, New Zealand) using a QIAGEN DNeasy Blood & Tissue Kit following the manufacturer’s protocol for tissue.

Template DNA was blunt-end repaired and had custom adapters ligated following the library preparation protocol of Meyer and Kircher (2010); this protocol was performed at the dedicated ancient DNA facility at the Australian Centre for Ancient DNA, University of Adelaide (Adelaide, Australia). Each adapter (5’ and 3’) contained a unique 7 mer index, allowing for identification of libraries and downstream removal of contaminant sequencing reads. Following library preparation, each sample was amplified using PCR in eight separate reactions to reduce amplification bias. Each 25 μL reaction contained 1 × PCR buffer, 2.5 mM MgCl_2_, 1 mM dNTPs, 0.5 mM primer, 1.25 U AmpliTaq Gold, 2 μL DNA extract. Reactions were subjected to the following thermocycling regime: 94°C for 12 min; 13 cycles of 94°C for 30 s, 60°C for 30 s, 72°C for 40 s (plus an additional 2 s per cycle); and a final extension of 72°C for 10 min. Individual PCR products for each library were pooled following amplification and purified using AMPure magnetic beads (Agencourt).

Hybridisation enrichment was conducted using the same baits as for the Brazilian merganser. Enrichment was performed using 200 ng of library following the manufacturer’s recommended protocol (myBaits v1), with the exception that the incubation was extended to 44 h (3 h at 60°C, 12 h at 55°C, 12 h at 50°C, 17 h at 55°C). Following incubation, baits were immobilised on magnetic MyOne Streptavidin Beads (Life Technologies, New Zealand). The baits were washed once with 1 × SCC and 0.1% SDS (15 min at room temperature), and twice with 0.1 × SCC and 0.1% SDS (10 min at 50°C), then resuspended in 0.1 M NaOH pH 13.0. The resulting enriched library was purified using a MinElute spin-column (QIAGEN) and subjected to a further round of PCR (12 cycles, eight reactions, using above recipe). Enriched libraries were subjected to a final round of PCR (7 cycles, 5 x 25 μL reactions, using above recipe) with fusion primers to add full-length Illumina sequencing adapters. Libraries were diluted to 2 nM and sequenced on an Illumina MiSeq using 2 × 150 (300 cycle, paired-end) sequencing chemistry at the School of Agriculture, Food, and Wine, University of Adelaide (Adelaide, Australia).

### Data processing

Raw sequencing reads for the Brazilian merganser library were demultiplexed using BaseSpace based on its i5 and i7 index sequences, while sequencing reads from the Auckland Island merganser were demultiplexed using “sabre” (http://github.com/najoshi/sabre) according to their unique 7-mer barcode combinations. For data from both libraries, we used AdapterRemoval v2.1.2 (Schubert *et al*., 2016) to trim residual adapters and low-quality bases (<Phred20 –minquality 4); merge overlapping paired-end reads (minimum overlap = 11 bp); and discard merged reads <30 bp (–minlength 30). Read quality was visualised using fastQC v0.10.1 (https://www.bioinformatics.babraham.ac.uk/projects/fastqc/) before and after trimming to ensure the trimming was efficient.

We mapped 1,480,994 merged reads from the Brazilian merganser and 377,348 merged reads from the Auckland Island merganser to the published mitogenome sequence of the common merganser (accession number MW849284) using BWA v.0.7.8 (Li & Durbin, 2009) and the backtrack (aln) algorithm with commonly used parameters for ancient DNA (i.e., -l 1024, -n 0.01, -o 2). Reads with mapping quality lower than a Phred score 25 were removed using SAMtools v.1.4 (Li *et al*., 2009). Duplicates were filtered using FilterUniqueSAMCons.py (Kircher, 2012). Mapped reads were visualised using Geneious v.9.1.6 (Biomatters, http://www.geneious.com, Kearse *et al*., 2012) and a majority-rule consensus sequence was created for each species. We then used these consensus sequences as the references for a new round of mapping, after which an updated consensus sequence was generated for each species. We repeated this process iteratively until no more reads mapped to the reference. A final 75% majority consensus sequence was then generated for each library and checked by eye in Geneious, calling nucleotides only for positions with a minimum depth-of-coverage of 3x. Rates of deamination in the mapped reads from each library were assessed using AuthentiCT v.1.0.1 (Peyrégne & Peter, 2020) to ensure they exhibited patterns consistent with ancient DNA damage (i.e., an elevated frequency of C to T transitions at the beginning of reads).

### Phylogenetic analysis

We aligned our new mitogenomes from the Auckland Island merganser and Brazilian merganser with previously published mitogenomes of 17 other members of Mergini— including the remaining three extant species of *Mergus*—and one outgroup. Multiple sequence alignment was performed using the MUSCLE algorithm (Edgar, 2004) as implemented in Geneious. We extracted and concatenated sites from our alignment that belonged to four subsets: first, second, and third positions of H-strand protein coding genes, and rRNA coding genes. Ambiguously aligned columns were removed from the rRNA gene alignment using Gblocks v0.91b (Castresana, 2000) with default settings.

We inferred a maximum likelihood phylogeny using IQ-TREE v1.6.11 (Nguyen *et al*., 2015). Our analysis in IQ-TREE first involved using ModelFinder (Kalyaanamoorthy *et al*., 2017) to determine the best partitioning scheme and nucleotide substitution models for each subset of sites (according to the Bayesian Information Criterion): K3Pu+F+R2 for a partition comprising first codon positions and rRNA genes; TIM3+F+R2 for a partition comprising second codon positions, and TIM3+F+I+G4 for a partition comprising third codon positions. Each partition had its own evolutionary rate, but all partitions contributed to the branch lengths of a single best tree (-spp; Chernomor *et al*., 2016). We assessed topological support using 1000 ultrafast bootstrap replicates (resampling within partitions; Hoang *et al*., 2017). Finally, we tested the tree with the highest likelihood against several alternative topologies (i.e., different relationships between *Mergus* species, including a sister relationship between the Auckland Island merganser and Brazilian merganser) using the KH (Kishino & Hasegawa, 1989), SH (Shimodaira & Hasegawa, 1999), and AU tests (Shimodaira, 2002) as implemented in IQ-TREE.

We estimated a time-scaled Bayesian phylogeny using BEAST v1.8.4 (Drummond & Rambaut, 2007). First, we removed the outgroup sequence from our alignment, then ran PartitionFinder v1.1.1 (Lanfear *et al*., 2012) on the same four subsets of alignment sites. In this case the best partitioning scheme and nucleotide substitution models (according to the Bayesian Information Criterion) was: K81uf+I+G for a partition comprising first codon positions and rRNA genes; TrN+I+G for a partition comprising second codon positions, and GTR+I+G for a partition comprising third codon positions. Substitution rate was modelled with a single lognormal relaxed clock (including a rate multiplier parameter for each partition), and we applied a Birth-Death tree prior.

To calibrate the substitution rate, we constrained the time to most recent common ancestor of true mergansers (*Mergus* spp.) and the hooded merganser (*L. cucullatus*) according to a lognormal distribution such that 95% of the prior probability fell between 14 and 23 million years before present (mean = 4.48; standard deviation = 0.5; offset = 14.0). The lower bound (minimum age) of the prior distribution is based on the fossil merganser *Mergus miscellus* Alvarez and Olsen, 1978 from North America (Alvarez & Olsen, 1978), while the upper bound (maximum age) is based on an apparent lack of any fossil representatives of crown Mergini from the Oligocene. This timescale is also consistent with the Miocene smew (*Mergellus* spp.) and goldeneye (*Bucephala* spp.) fossils from Hungary (Gal *et al*., 1998) and previous divergence dates for scaly-sided merganser and common merganser in the mid-late Miocene (Sun *et al*., 2017).

We ran three independent BEAST Markov chain Monte Carlo chains; each chain was run for 10,000,000 generations, sampling every 1,000 generations. Convergence of parameter values was monitored using Tracer v1.7.2 (Rambaut *et al*., 2018) to ensure that effective sample sizes for all parameters were >200. The first 10% of sampled trees of each chain were discarded as burn-in and the remaining trees were combined using LogCombiner (part of the BEAST software package) and summarised as a maximum clade credibility tree using TreeAnnotator (part of the BEAST software package).

## RESULTS

### Ancient mitochondrial genomes

Mitogenomes were recovered from the Auckland Island merganser and Brazilian merganser specimens (15,678 bp and 16,218 bp, respectively). For the Auckland Island merganser, the mean length of unique mapped reads (n = 35,562) was 66.9 bp (standard deviation = 24.1) and the mean depth-of-coverage was 151.6× (standard deviation = 60.4); for the Brazilian merganser the mean length of unique mapped reads (n = 206,710) was 69.8 bp (standard deviation = 25.3) and the mean depth-of-coverage was 921.3× (standard deviation = 315.3). Deamination rates at the terminal base of mapped reads (measured as the frequency of C-T mismatches at the first sequenced base) were 3.6% for the Auckland Island merganser and 2.3% for the Brazilian merganser, before steadily decreasing to <1% at positions further from the terminus. These patterns of deamination, and the short length of mapped reads, are consistent with patterns of damage expected in DNA extracted from ancient specimens.

### Phylogenetic analysis

Bayesian and Maximum Likelihood phylogenetic analyses produced highly concordant and moderately well-resolved phylogenies for Mergini, with most branches receiving strong bootstrap (BS) or Bayesian posterior probability (PP) support. The main exception is the phylogenetic placement of the harlequin duck (*Histrionicus histrionicus*). In our Maximum Likelihood analysis the harlequin duck was sister to a clade comprising scoters (*Melanitta* spp.), goldeneyes, smew, hooded merganser, and true mergansers (*Mergus* spp.) with 95% BS. In contrast, in our Bayesian analysis, the harlequin duck was sister to a clade comprising the long-tailed duck (*Clangula hyemalis*) and the eiders (*Polysticta stelleri* and *Somateria* spp.) with 0.78 PP.

Bayesian and Maximum Likelihood phylogenetic analyses both resulted in a strongly supported clade comprising smew, hooded merganser, and *Mergus* (98 BS, 1.0 PP), which was sister to goldeneyes (100 BS, 0.97 PP). Sequences from living and extinct *Mergus* species formed a strongly supported monophyletic clade (100% BS, 1.0 PP), within which the red-breasted merganser was sister to a strongly supported clade comprising the scaley-sided merganser, Brazilian merganser, common merganser, and Auckland Island merganser (97% BS, 1.0 PP). While the Auckland Island merganser appears to be the sister species to the common merganser, this relationship was only weakly supported (73% BS, 0.85 PP)—indeed, an alternative tree where the Auckland Island merganser and Brazilian merganser were sister species could not be rejected at the p = 0.05 level using the KH, SH, and AU tests as implemented in IQ-TREE.

Our time-calibrated phylogeny indicates the common ancestors of most extant Mergini genera occurred during the late Oligocene or early Miocene. The mean estimate for the common ancestor of crown *Mergus* was 13.7 million years ago (95% HPD = 11.0 – 17.5 million years ago). The most recent node within *Mergus* was the common ancestor of the Auckland Island merganser and common merganser for which we estimated a mean age of 9.1 million years (95% HPD = 7.0 – 12.0 million years ago).

## DISCUSSION

### Merganser biogeography and the origin of the Auckland Island merganser

Our phylogenetic analysis was generally well-resolved and broadly concordant with previous studies of Mergini, taking into account the absence of previously un-sequenced taxa that may change some evolutionary relationships (Livezey, 1995; Gonzalez *et al*. 2009; Solovyeva & Pearce, 2011; Liu *et al*. 2012; Sun *et al*. 2017; Buckner *et al*. 2018; Lavretsky *et al*. 2021).

The tribe Mergini are almost entirely restricted to the Northern Hemisphere, with some species exhibiting widespread distributions, while others have more restricted distributions in Eurasia or North America. It is likely that the ancestral ‘merganser’ (*Mergellus, Lophodytes*, and *Mergus*) had a Northern Hemisphere distribution, was migratory in order to access seasonal riverine and marine habitats, and nested in cavities (Kear, 2005; Williams *et al*., 2012; Williams *et al*., 2014).

The ‘mergansers’ split from the North American goldeneyes around 25 million years ago (95% HPD 21.9 – 34.6 million years ago), pre-dating mean estimates for the divergences of the smew, hooded merganser, and *Mergus* by over four million years later (Figure 4). A 12-13 million year old fossil assigned to the smew lineage is consistent with our divergence datesand the presence of smew in Eurasia since at least the middle Miocene (Gal *et al*., 1998). The hooded merganser, which is restricted to North America, diverged from *Mergus* around 18 million years ago (95% HPD = 15.0 – 22.0 million years ago) and is consistent with the 14 million year old Miocene *M. miscellus* fossil from North America (Alvarez & Olsen, 1978). Within *Mergus*, the red-breasted merganser, which has a widespread Northern Hemisphere distribution, diverged about 14 million years ago (95% HPD = 11.1 – 17.6 million years ago). This was followed by the divergence of the scaly-sided merganser of northeast Asia, the Brazilian merganser, the Auckland Island merganser, and the widespread Northern Hemisphere common merganser (mean node ages between 9.2 and 11.1 million years ago). The deep divergences within *Mergus* are in stark contrast to more recent divergences between sister species within goldeneyes and eiders, which have mean ages no older than 2.4 million years (Figure 4).

**Figure 1.**
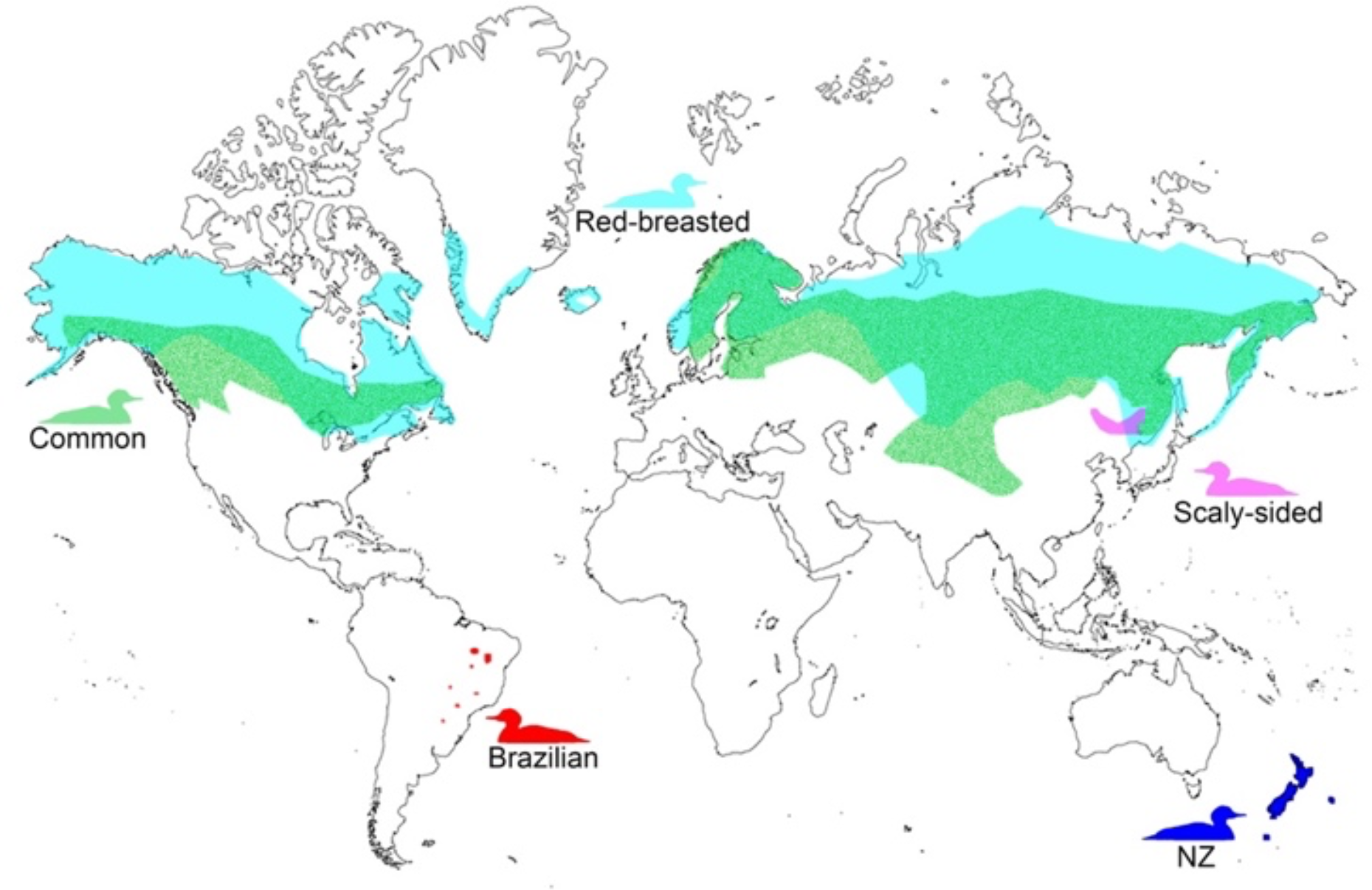
Schematic of the breeding distributions of *Mergus* spp. The New Zealand (NZ) radiation encompasses the Auckland Island merganser (465 km south of NZ) and Chatham Island merganser (785 km east of NZ), as well as *Mergus* spp. from mainland NZ. The distribution of the hooded merganser (*Lophodytes cucullatus*), which is restricted to North America, is not shown. Breeding distributions are based off the Cornell Lab of Ornithology Birds of the World website.

**Figure 2.**
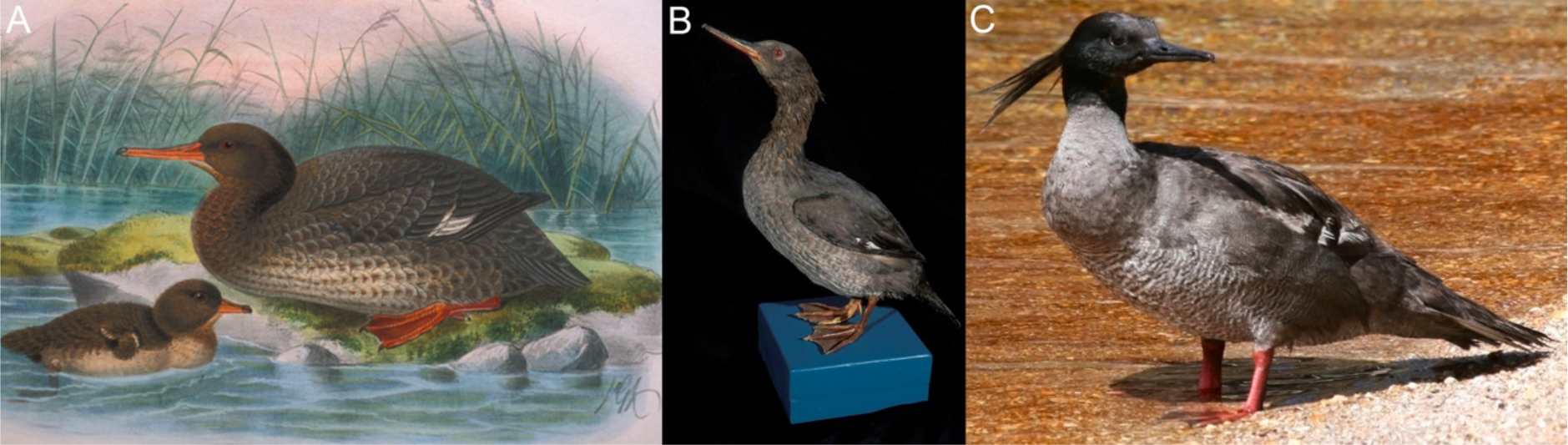
In the Southern Hemisphere mergansers are only known from the New Zealand region and South America, represented here by the Auckland Island merganser (A, artistic reconstruction by JG Keulemans from Buller (1888); B, historical museum skin, Otago Museum A51.50, photo by Rod Morris), and (C) the Brazilian merganser (photo by Savio Freire Bruno CC BY-SA 3.0).

**Figure 3.**
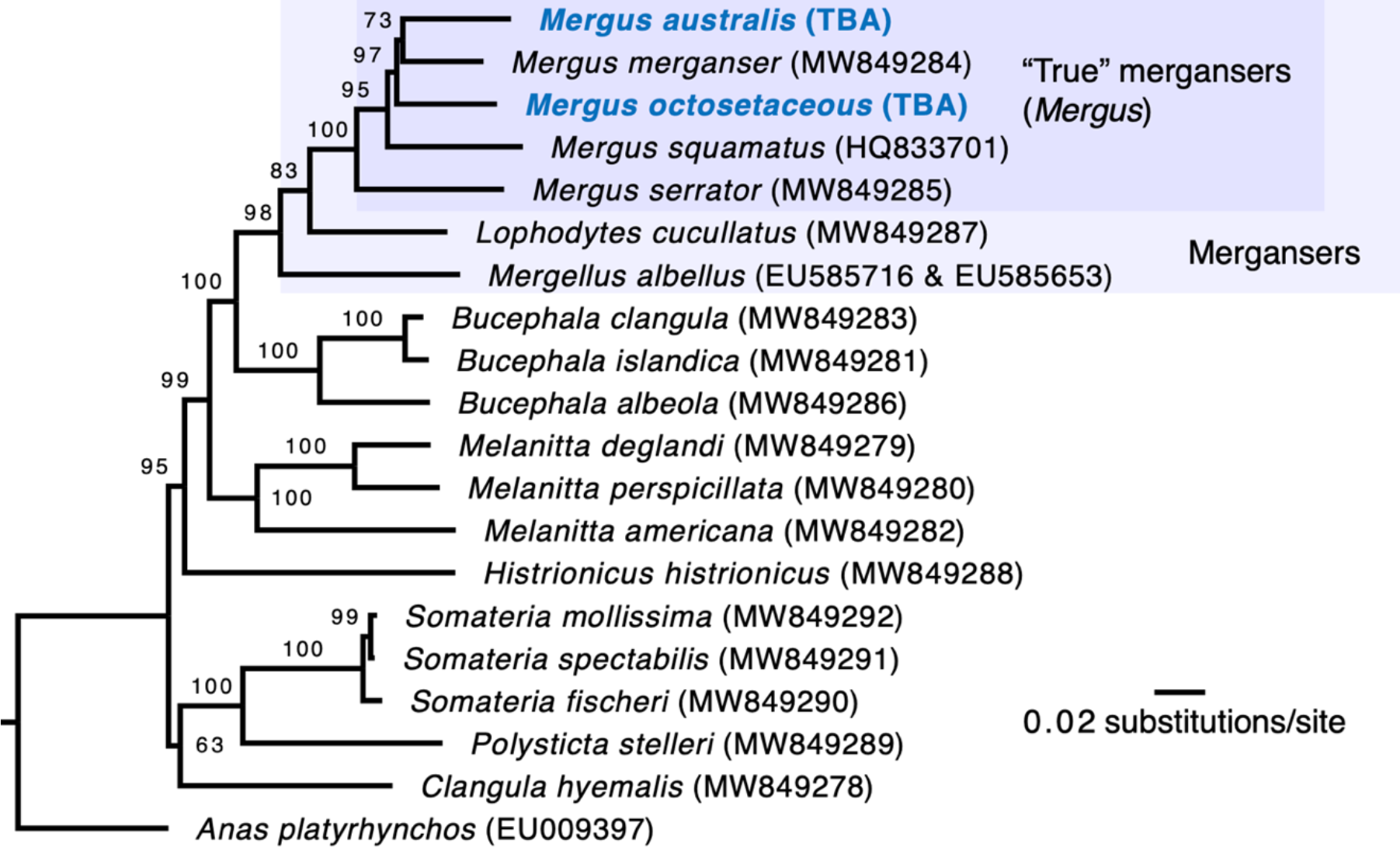
Maximum Likelihood phylogeny of Mergini mtDNA created using IQ-TREE. Branch lengths are proportional to the number of substitutions. Bootstrap support values are shown above branches. Tips are labelled with species names and GenBank accession numbers.

**Figure 4.**
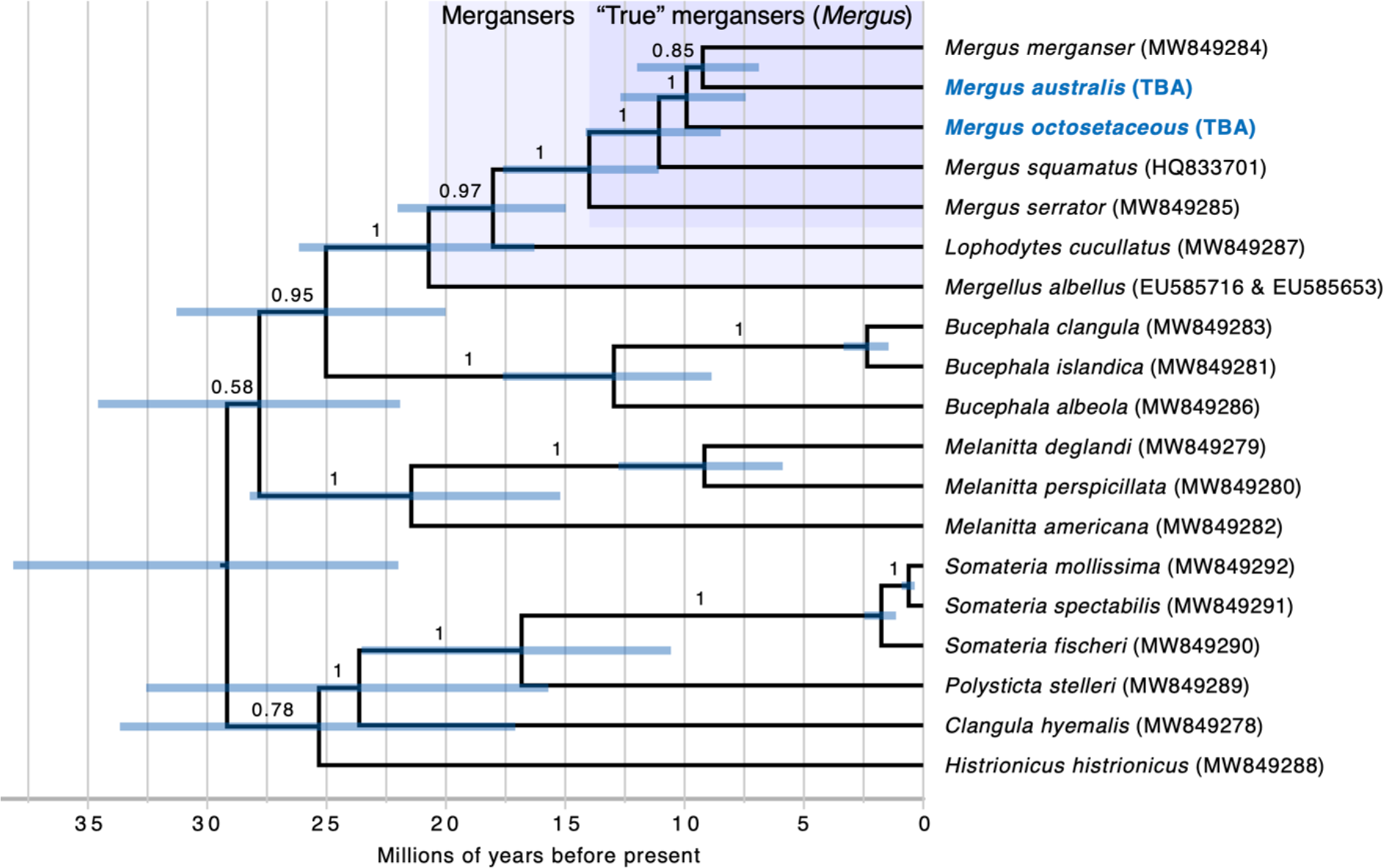
Time-calibrated Bayesian phylogeny of Mergini mtDNA created using BEAST. Branch lengths are proportional to time in millions of years before present. Posterior probability support values are shown above branches. Blue bars represent the 95% highest posterior densities for node age estimates. Tips are labelled with species names and GenBank accession numbers.

The low support for the sister group relationship between the Auckland Island merganser and common merganser (0.85 PP, 73% BS) is likely due to a lack of phylogenetic signal, with the time between divergence events being too short to accumulate enough informative mutations. Therefore, the phylogenetic relationship between the Brazilian merganser, Auckland Island merganser, and common merganser should be interpreted as an unresolved polytomy. Despite previous morphology based phylogenetic analyses suggesting the Brazilian merganser and Auckland Island merganser were the result of separate colonisation events (Livezey 1989, 1995), we cannot reject the possibility that they are sister taxa (in line with some behavioural and plumage characters; Johnsgard, 1961) originating from a single colonisation event of the Southern Hemisphere at least seven million years ago. Further palaeogenomic data incorporating nuclear DNA (e.g. McCormack *et al*., 2016) may help resolve the relationships between living and extinct mergansers.

Given the Brazilian merganser and Auckland Island merganser are nested within Northern Hemisphere *Mergus* congeners, it is likely that these lineages originated from northern migrants. All Northern Hemisphere mergansers are migratory to varying degrees (Livezey, 1995; Kear, 2005). The specific habitats occupied by these southern species may have negated the need for migration between winter and summer, and breeding and non-breeding grounds. It remains possible that further extinct Mergini taxa will be discovered as scientists increasingly focus on Southern Hemisphere terrestrial and marine fossil localities (e.g. Tennyson & Mannering, 2018; Tennyson *et al*., 2022; Tennyson & Salvador, 2023).

Evolutionary genetic research into the geographic origin and arrival times of the ancestors of New Zealand birds has largely neglected anatids. Mitchell *et al*. (2014b) dated the arrival of teals (*Anas* spp.) to 2.85 million years ago (95% HPD 2.85 – 1.78 million years ago), compared to our conservative estimate of at least seven million years ago for mergansers. Further research is needed to determine how changes in habitat preferences and niche availability over time has influenced the timing of arrival and diversification of anatids in New Zealand (e.g. Rawlence *et al*. 2019). Indeed, the riverine and seasonally coastal (i.e. in sheltered bays in winter) habitat of mergansers (Kear, 2005; Williams *et al*. 2012; Williams *et al*. 2014), and the presence of subfossil and archaeological bones in coastal sand dune deposits throughout the New Zealand region (including taxonomically indeterminate mainland specimens *Mergus* spp.) represents an ideal opportunity to further test for phylogeographic structure and speciation events.

Among anatids, the riverine (and potentially coastal; Buller, 1888) blue duck (*Hymenolaimus malacorhynchos*) can be thought of as a broad ecological analogue of mergansers, albiet at a lower trophic level (Collier & Lyon, 1991). The blue duck exhibits strong phylogeographic structure across Cook Strait (that separates the North and South Islands) that likely dates to the late Pleistocene (Grosser *et al*., 2016), potentially coincident with formation of the strait around 500,000 years ago (Trewick & Bland, 2012). Earlier divergence dates, coincident with the narrowing of the Manawatu Strait 5 – 1.5 million years ago, which separated the South Island and lower North Island from the upper North Island (Trewick & Bland, 2012), have been seen in other birds (e.g. Lubbe *et al*., 2022). If the Chatham Islands merganser (Williams *et al*., 2014) is genetically distinct from mainland mergansers, it should date to no older than the emergence of the islands three million years ago (Campbell, 2008), as observed for other endemic lineages (McCulloch & Waters, 2019 and references therein). Likewise, the Auckland Islands emerged during the early to middle Miocene conservatively at least 12 million years ago but they were heavily glaciated during the Pleistocene (Scott & Turnbull, 2019). Colonisation of the Auckland Islands, likely from mainland New Zealand (e.g. Mitchell *et al*., 2014b; Rawlence *et al*., 2015), could have occurred any time since the arrival of mergansers in the New Zealand region 7 – 12 million years ago, but potentially not until the late Pleistocene given the likely impact of glaciation on available habitat.

## CONCLUSION

Our analyses have shown that mergansers colonised the Southern Hemisphere at least seven million years ago during the late Miocene. However, we were unable to resolve whether there were two separate colonisation events, or a single dispersal pulse followed by subsequent divergence into the Brazilian merganser and Auckland Island merganser. Nuclear (palaeo)genomic data may help to further resolve the relationship between living and extinct mergansers, including other merganser fossils from New Zealand that have not yet been subjected to palaeogenetic analysis, and how divergence dates in mergansers compare to other anatids in the region across a range of diverse habitat types.

## ACKNOWLEGEMENTS

We thank Canterbury Museum (Paul Scofield) and the American Museum of Natural History (Thomas Thrombone) for permission to sample their Auckland Island merganser and Brazilian merganser historical museum specimens, respectively. Thank you to Pascale Lubbe for helpful discussion on the biogeography of New Zealand birds. Māori are kaitiaki (guardians) of the fauna and flora of Aotearoa New Zealand, with which they are interconnected through shared whakapapa (genealogy) — genetic data from the miuweka (Auckland Island merganser) are published with the support of the Ngāi Tahu Murihiku Kaitiaki Rōpū Committee.

## FUNDING

Funding was provided by the Royal Society Te Apārangi Marsden Fund (16-UOO-096; 20-UOO-130) and the University of Otago.

## DATA AVAILABILITY

The data that support the findings of this study are available from … [DOIs and/or accession numbers TBA].

## Notes

### Competing Interest Statement

The authors have declared no competing interest.

